# United States stakeholder insights on genetic testing for equine health and breeding

**DOI:** 10.1101/2025.07.29.667538

**Authors:** Michael J. Mienaltowski, Sarah Hernandez, Emma Nastrini, Carissa L. Wickens, Molly E. McCue, Laura Patterson Rosa, Elaine M. Norton, Annette M. McCoy, Samantha A. Brooks

## Abstract

The S1094 United States Department of Agriculture (USDA) Multistate Research Project is a community of scientists working together to utilize equine genetics and genomics to improve the health and well-being of horses while also providing educational and genetic testing resources to those in the equine industry. To assess the knowledge and priorities of stakeholders regarding genetic testing, we conducted an online survey targeting horse owners and enthusiasts in the United States. Data were collected from March to October 2024 with 412 respondents from 44 states completing the survey. Most of the survey participants were horse owners, and the two most common age ranges for respondents were 31-50 (33.6%) and 51-70 (33.3%) years old. The two most common use categories for horses reported by respondents were sport and stock horses. The survey revealed that a majority of respondents were confident in their knowledge of genetics and that they relied primarily on breed and discipline organizations, as well as universities for their genetic information. Secondarily they relied upon social media and their veterinarians. Respondents expressed significant interest in genetic testing for diversity and performance traits. Across all horse use categories, arthritis, colic and intestinal disease, laminitis, metabolic disorders, and tendon and ligament issues had the greatest level of concern - all of which are complex genetic traits and will require collaborative efforts to discern levels of genetic involvement. Results from the survey underscore the need for continued outreach and education of horse owners, as well as further development of genetic testing tools to support informed decision-making among stakeholders in the equine industry.

**Lay Summary:** Genetic testing helps horse owners, breeders, and breed registries and associations verify parentage, evaluate health, and improve performance. The S1094 USDA Multistate Research Project has a mission of improving genomics and genetics tools, as well as providing educational resources for those in the equine industry. To better understand the knowledge and priorities of stakeholders in the equine industry, we conducted an online survey netting responses from 412 individuals in the United States. Respondents were primarily horse owners who self-identified as confident in their knowledge of horse genetics. Furthermore, those responding indicated that their primary resources for information about genetics were breed and discipline association websites and universities. They also expressed interest in genetic testing for diversity and performance traits. Across all horse use categories, arthritis, colic and intestinal disease, laminitis, metabolic disorders, and tendon and ligament issues had the greatest level of concern. Given the complex nature of these conditions, collaborative efforts will be required to discern levels of genetic involvement. Respondents felt that genetic testing and education efforts have significant impacts on the equine industry because they aid owners in making breeding decisions and in the management of horse health and well-being.

**Teaser Text:** This nationwide survey of U.S. horse owners discerned strong stakeholder interest in equine genetic testing, determined sources for knowledge acquisition, and highlighted key priorities for the development of genetic tools and educational materials.

## Introduction

The equine industry in the United States comprises many small and distinct enterprises with no major entities or companies dominating the market. Collectively, though, the equine industry is quite large with an estimated total impact of $177 billion annually (American Horse Council Foundation, 2023). A wide variety of businesses and services working directly and indirectly within the equine industry contribute 2.2 million jobs to the United States economy; these include breed registries, horse production and management, venues for sports and activities, veterinary services, and other such allied businesses (American Horse Council Foundation, 2023). In order to maintain a robust industry, equine producers must provide high quality horses for work, sport, entertainment, and companionship. Thus, understanding priorities, regional or sub-sectional differences and stakeholder engagement are essential to supporting the sustainable growth and productivity of the equine industry.

Horse owners and breed registries work together to maintain breed standards in the industry. From the very founding of a breed, producers have made efforts to identify, verify parentage, and evaluate their horses (Brooks, 2021). Genetic testing is a tool to both track lineage and evaluate health and potential performance of horses in breeding populations (Bellone & Avila, 2020). Breed registries require genetic testing for horses to become members of the breed associations, with membership having implications on level of participation within the industry and marketable value of the animal. Over the years, genetic testing has also been used to identify diseases or problematic traits so that owners and breed organizations can minimize or manage the incidence of those undesirable traits. Hence, utilization of genetic testing is a powerful tool to improve the welfare and performance of horses.

Industry members working directly with horses generally have substantial practical knowledge related to performance-limiting conditions and diseases in their breed or discipline of interest. While horse breeding is historically as much art as science, the use of genetic testing is becoming much more prevalent as owners and breeders work to optimize matings as well as to learn more about their horses to better tailor their management and predict their performance. However, more can be done to reach horse owners and breeders. A study based within the state of Florida found that familiarity with genetic testing was one barrier to genetic testing (Hammons, et al. 2020). Thus, it is essential that stakeholders in the horse industry are aware of genetic testing and its appropriate applications. Moreover, because stakeholders are interested in the improvement of their horses, scientists have an opportunity both to reach stakeholders to provide knowledge about existing genetic testing options and to learn more about stakeholder priorities for genetic improvement. For decades, scientists at institutions across the United States have worked together within an equine genetics and genomics community focused on improving the health and well-being of horses. They have come together to develop genetic tests for specific diseases affecting horses or beneficial traits, to examine breed differences, and even to address population structure. The reader may refer to any of the several studies and reviews for examples (Avila, et al. 2018; Hill, et al. 2019; Lindgren, et al. 2020; Tallmadge, et al. 2020; Aleman, et al. 2022; Bailey, et al 2022; Kingsley, et al. 2023; Durward-Akhurst, et al. 2024; McFadden, et al. 2024; Finno, 2025).

In 2022, a USDA multistate project entitled, “S1094: Genomic tools to improve equine health, wellbeing and performance” was created to support this collaborative network. One mission of the multistate project is to expand the availability of genetic diagnostic testing and education on its use to equine industry stakeholders. To reach stakeholders, discern their understanding of genetics, and seek out their opinions on what they consider their performance and disease trait priorities, we developed a needs assessment and distributed it to stakeholders in the spring and summer of 2024. This is a summarization of the findings from that assessment.

## Materials and Methods

A survey was conducted using Qualtrics XM (Qualtrics, LLC, Seattle, WA), a cloud-based survey platform. This survey was reviewed by the Internal Review Board (IRB) Office at the University of Florida and an IRB exemption was provided (#ET00023502). The targeted survey demographic was horse owners and enthusiasts in the United States who were at least 18 years of age. The survey was distributed to stakeholders via email lists as a part of university, breed and professional association list-servs, as well as broadly through social media channels like Facebook (Meta Platforms). Data were collected from 2024-03-06 until 2024-10-15.

The survey comprised 38 questions. At the start of the survey there were nine questions to inquire about the demographics reached by the survey. These inquiries included: owner age, location (country and United States ZIP code), respondent’s primary role in the equine industry, type of horse focus (breed type and/or discipline), numbers of horses owned or cared for, type of riding activity that was respondent’s main focus, level of genetics knowledge, and preferred sources for receiving information about horse genetics. From a respondent’s answer to “type of horse focus,” for certain responses, they were directed to questions relevant to their focus breed type/discipline: in blocks of questions: racing (thoroughbreds, standardbreds, and quarter horses), sport horse (warmbloods, Friesians, Andalusians, and thoroughbreds), stock horse (APHA, AQHA, and APHC), Arabian, heavy and draft horses, miniature horse and ponies. Within these blocks, questions could include inquiries into specific breeds worked with, genetic conditions relevant to those breed types/disciplines concerning them most, an open-ended question on conditions that respondents suspect are inherited, interest in knowing level of genetic diversity, and interest in genetic testing for performance genes. Finally, they were directed to a question regarding their general concerns regarding horse health issues including arthritis, asthma or heaves, behavior or temperament, exercise induced pulmonary hemorrhage, cardiac disorders, colic or intestinal disease, eye disorders, laminitis, metabolic syndrome or Pituitary Pars Intermedia Dysfunction (PPID), muscle disorders, navicular syndrome, reproduction issues or embryonic loss, tendon or ligament issues, severe infectious diseases, laryngeal hemiplegia, ulcers, and Wobblers.

Survey results were organized and analyzed using Excel 2016 (Microsoft, Redmond, WA), Prism 10.4.1 (GraphPad, Boston, MA), and RStudio v.2024.12.1+563 (Posit, Boston, MA). Comparison of responses between respondents (grouped by number of horses owned or cared for), was done using the Mann-Whitney test, with significance set at p<0.05. Responses to free-response questions were analyzed using tools built into the Copilot Artificial Intelligence (AI) (Microsoft, Redmond, WA) program. Briefly, when respondents were asked about how genetics impacts horse business, operations, and activities, the prompt was open-ended thus allowing respondents to write as much or as little as they chose. Copilot AI was used to broadly categorize and tally responses for the prompt. Moreover, when respondents were asked about which conditions they felt might be heritable, Copilot AI was instructed to make text size proportional with the number of mentions in survey responses.

## Results

### General Respondent Demographics

A total of 412 respondents from the United States completed the survey. Respondents were from 44/50 states with Florida having the most responses **(Figure 1)**. Respondent ages varied with the two most common categories 31-50 years old (35.92%) and 51-70 years old (35.68%) **(Figure 2A)**. Of the respondents, 61.0% identified as owners, followed by 14.6% as breeders and 5.8% as veterinary practitioners **(Figure 2B)**. Most of the respondents (62.1%) indicated that they owned or cared for 1-5 horses, while 17.0% had 5-10 horses and 17.9% owned or cared for more than 10 horses **(Figure 2C)**. When asked about their major focus of horse use (categorized by breed type or discipline), the most frequent response was sport horses (33.0%), followed by stock horses (30.3%), Arabian horses (12.4%), gaited horses (10.0%), heavy and draft horses (6.3%), and racing horses (5.1%) **(Figure 2D)**.

**Figure 1.**
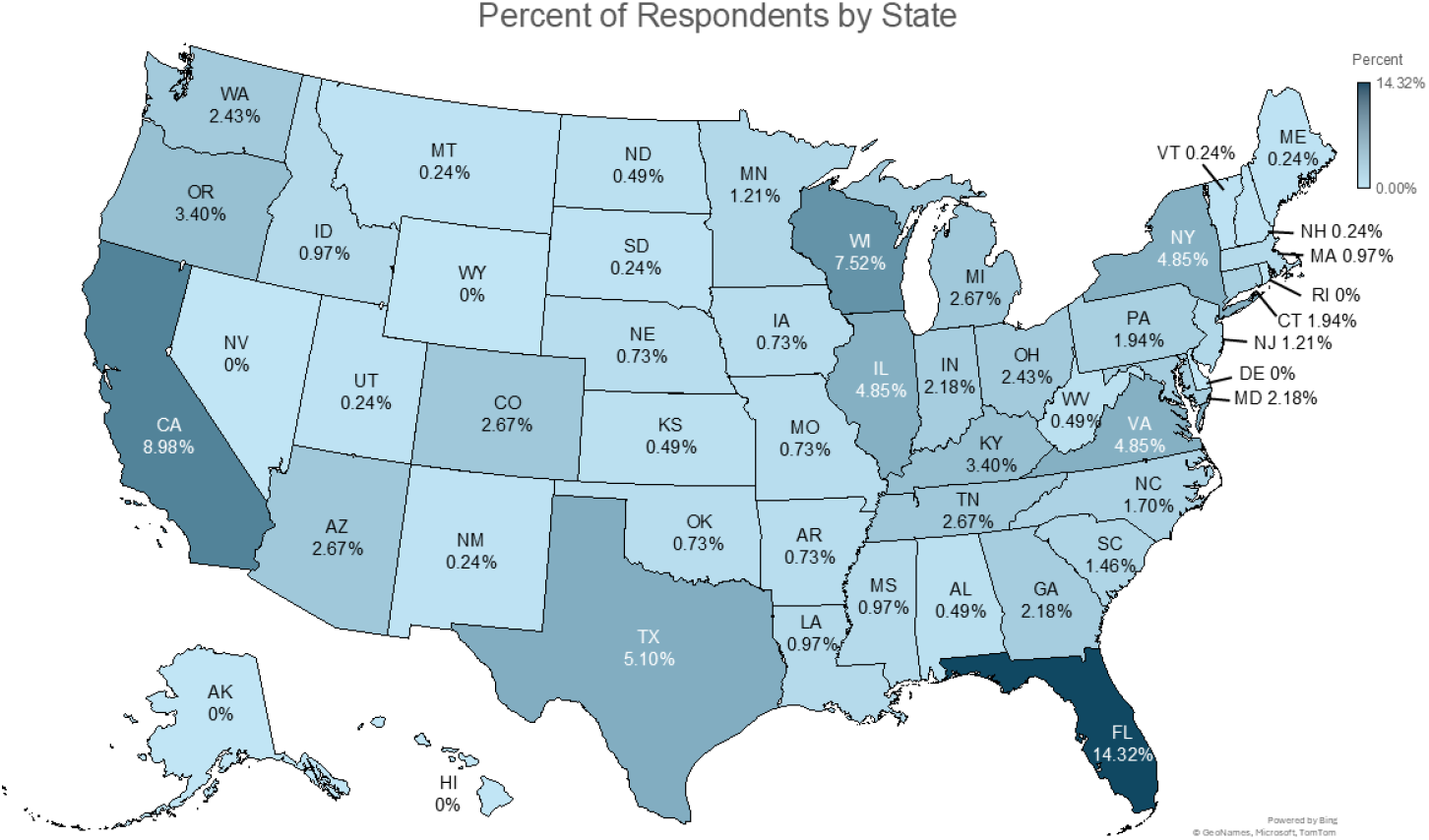
Distribution of Respondents. Respondents (n=412) were found throughout the United States with participation occurring in 44 states. Six states (AK, DE, HI, NV, RI, and WY) were not represented. States with a higher percentage of respondents are darker blue.

**Figure 2.**
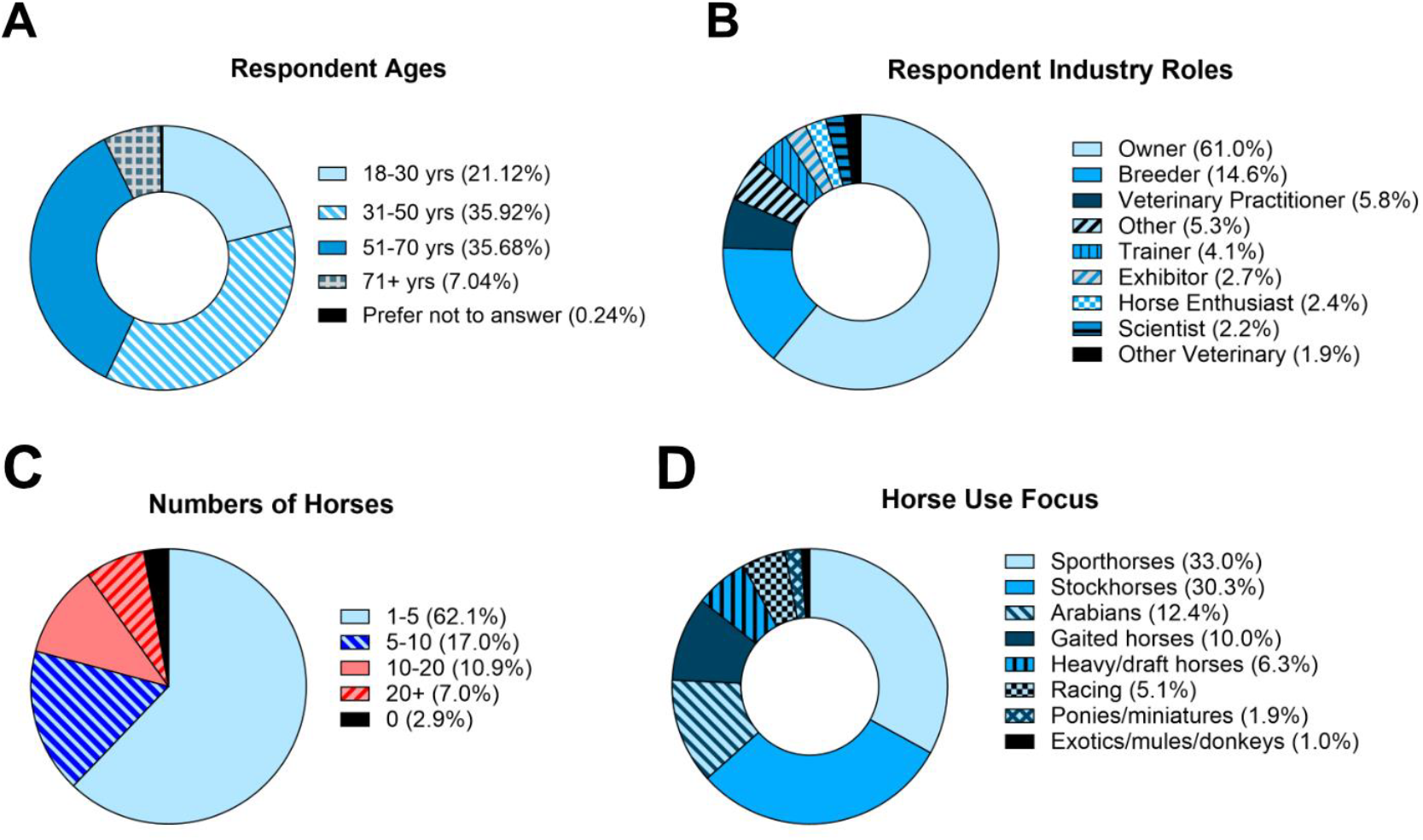
Survey Respondent Demographics. Survey findings included: **(A)** respondent ages, **(B)** industry roles, **(C)** numbers of horses cared for or owned, and **(D)** major focus of horse use amongst respondents.

Respondents were also asked about their knowledge and familiarity with genetics. When asked about their level of confidence in the knowledge of genetics using a four-point scale (scale: 1, little knowledge; 2, high school level science understanding; 3, confident; 4, expert knowledge), we saw a small but statistically significant difference in confidence depending upon the number of horses for which they cared or owned. A greater proportion of those with 10+ horses indicated having expert level and confident knowledge (12.2% and 77.0%, respectively) than did those with 0-10 horses (6.7% and 68.7%, respectively) (p=0.0045) **(Figure 3A)**. Respondents indicated that they most often relied on breed and discipline organizations as well as universities for their horse genetics information, though horse owner communities and affiliated social media, and veterinarians were also common sources of information **(Figure 3B)**.

**Figure 3.**
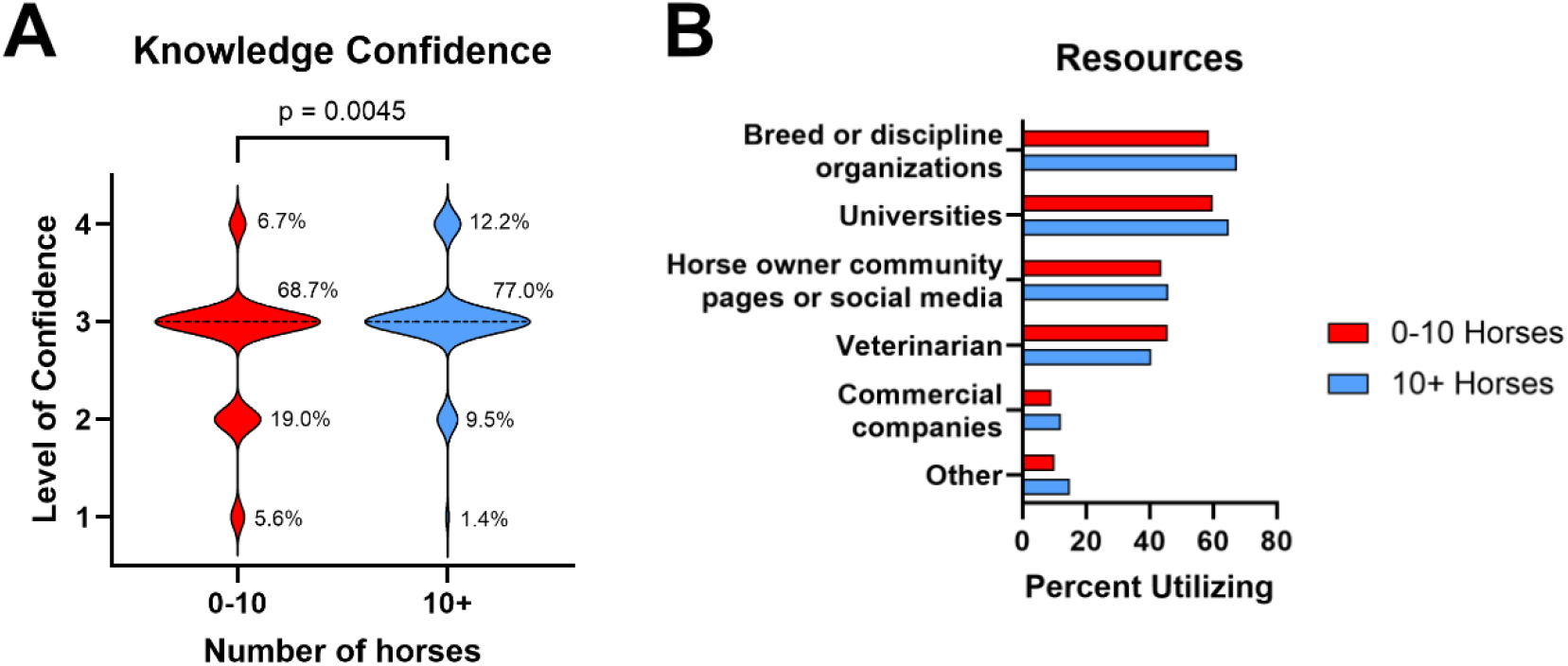
Respondent Knowledge and Resources Used. Respondents indicated **(A)** their level of confidence in the knowledge of genetics concepts (scale: 1, little knowledge; 2, high school level science understanding; 3, confident; 4, expert knowledge), as well as **(B)** which resources they use to obtain their knowledge. Comparisons were made between respondents caring for or owning 0-10 horses versus 10+ horses (Mann-Whitney test, p= 0.0045; 0-10 horses n = 268; 10+ horses, n = 74).

Respondents within three use disciplines were also asked about which known genetic conditions concerned them and about their interest in genetic testing. Within the category of sport horses, degenerative suspensory ligament desmitis (DSLD), osteochondritis dissecans (OCD), and cervical vertebral stenotic myelopathy (CVSM) were of greatest concern **(Figure 4A)**. For stock horses, polysaccharide storage myopathy (PSSM1), hyperkalemic periodic paralysis (HYPP), and hereditary equine regional dermal asthenia (HERDA) were of greatest concern **(Figure 4B)**. Racing horse owners shared OCD concerns with sport horse owners and HYPP with stock horse owners **(Figure 4C)**. Most respondents in these three groups indicated that they were interested in testing for genetic diversity, with 71.1%, 69.1%, and 52.4% owning sport horses, stock horses, and racing horses, respectively, answering affirmatively **(Figure 5A)**. There was also interest in testing for performance genes with 66.7%, 54.8%, and 53.8% of owners with racing horses, stock horses, and sport horses, respectively, indicating that they would like to see tests developed in this space **(Figure 5B)**.

**Figure 4.**
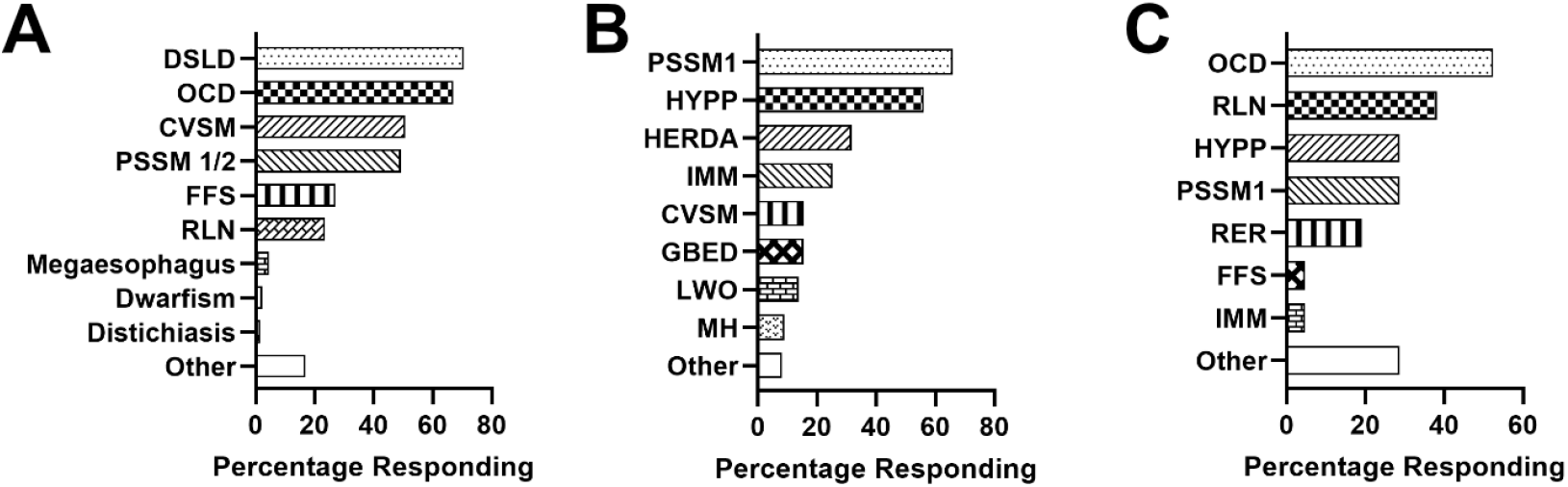
Genetic Conditions of Concern to Respondents. Respondents who owned or cared for three categories of horses were asked about which conditions with a known genetic component concerned them. Owners could provide multiple answers. Responses were given for (A) Sport horses, (B) Stock horses, and (C) Racing horses.

**Figure 5.**
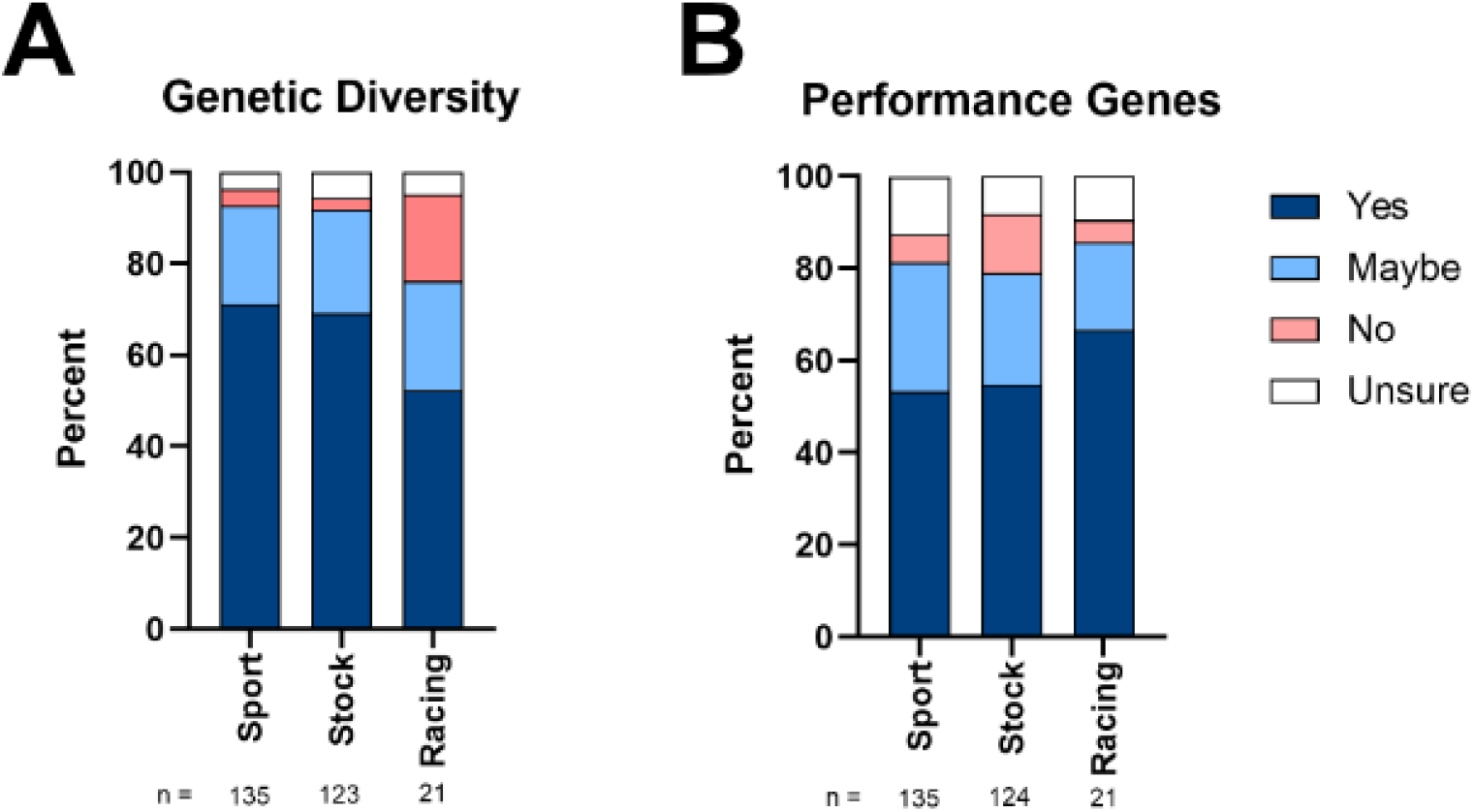
Interest in Testing for Genetic Diversity and Performance Genes. Sport, stock, and racing horse respondents were asked about their interest in testing for genetic diversity (A) and for performance genes (B).

All respondents were asked about how genetics impacts horse business, operations, and activities **(Table 1)**. Responses highlighted that genetics was important for breeding decisions (20%) as well as health and disease prevention (18%). Fifteen percent of respondents indicated that a horse’s performance and trainability could be impacted by genetics, and 12% responded that they felt that genetics had a role in a horse’s longevity and quality of life. Furthermore, 10% of responses indicated that there were economic implications to a horse’s genetic makeup. Finally, 8% of respondents indicated that knowledge of genetic testing helps owners make informed decisions and can lead to breed improvement.

**Table 1.** Broad themes of survey responses on “impact of genetics on horse business, operations, and activities.”

| Theme | Percent | Details |
| --- | --- | --- |
| Breeding Decisions | 20% | Importance of genetics in making informed breeding decisions to produce healthier and better-performing horses. |
| Health and Disease Prevention | 18% | Role of genetics in preventing and managing diseases, ensuring the overall health of horses. |
| Performance and Trainability | 15% | Genetics in relation to performance and trainability of horses, impacting their suitability for various activities. |
| Longevity and Quality of Life | 12% | Influence of genetics on the longevity and quality of life of horses, making them more resilient and healthier. |
| Economic Impact | 10% | Economic implications of genetics, such as affecting sales prices and reducing costs associated with health issues. |
| Genetic Testing and Knowledge | 8% | The importance of genetic testing and having knowledge about genetic predispositions as a key factor in making informed decisions. |
| Breed Improvement | 7% | The potential for genetics to improve breeds by eliminating undesirable traits and enhancing desirable ones. |

### Findings by Category of Horse Use

Once respondents indicated their primary horse use, they received questions specific to their use category. We will discuss findings from survey respondents responding with 5% or more by horse use category: sport horses, stock horses, Arabians, gaited horses, draft and heavy horses, and racing horses. Less than 2% of respondents provided answers for ponies and miniature horses and for exotic equids.

Those working with sport horses were the most common survey responders (33%). Respondents were asked about the breeds with which they were affiliated; most owned or cared for warmbloods (68.2%), followed by Thoroughbreds (19.4%) **(Table S1)**. When asked about the level of concern for various herd health issues, these respondents prioritized the following concerns: arthritis, behavior and temperament, colic and intestinal disease, laminitis, metabolic disorders, tendon and ligament issues, and ulcers **(Figure 6)**. When respondents were asked to identify other health issues of concern to their breeds/disciplines that they suspected of being heritable, the conditions provided most often by stakeholders were degenerative suspensory ligament desmitis (DSLD), overriding dorsal spinous processes (aka “kissing spines”), anhidrosis, and equine complex vertebral malformation (ECVM) **(Figure S2A)**.

**Figure 6.**
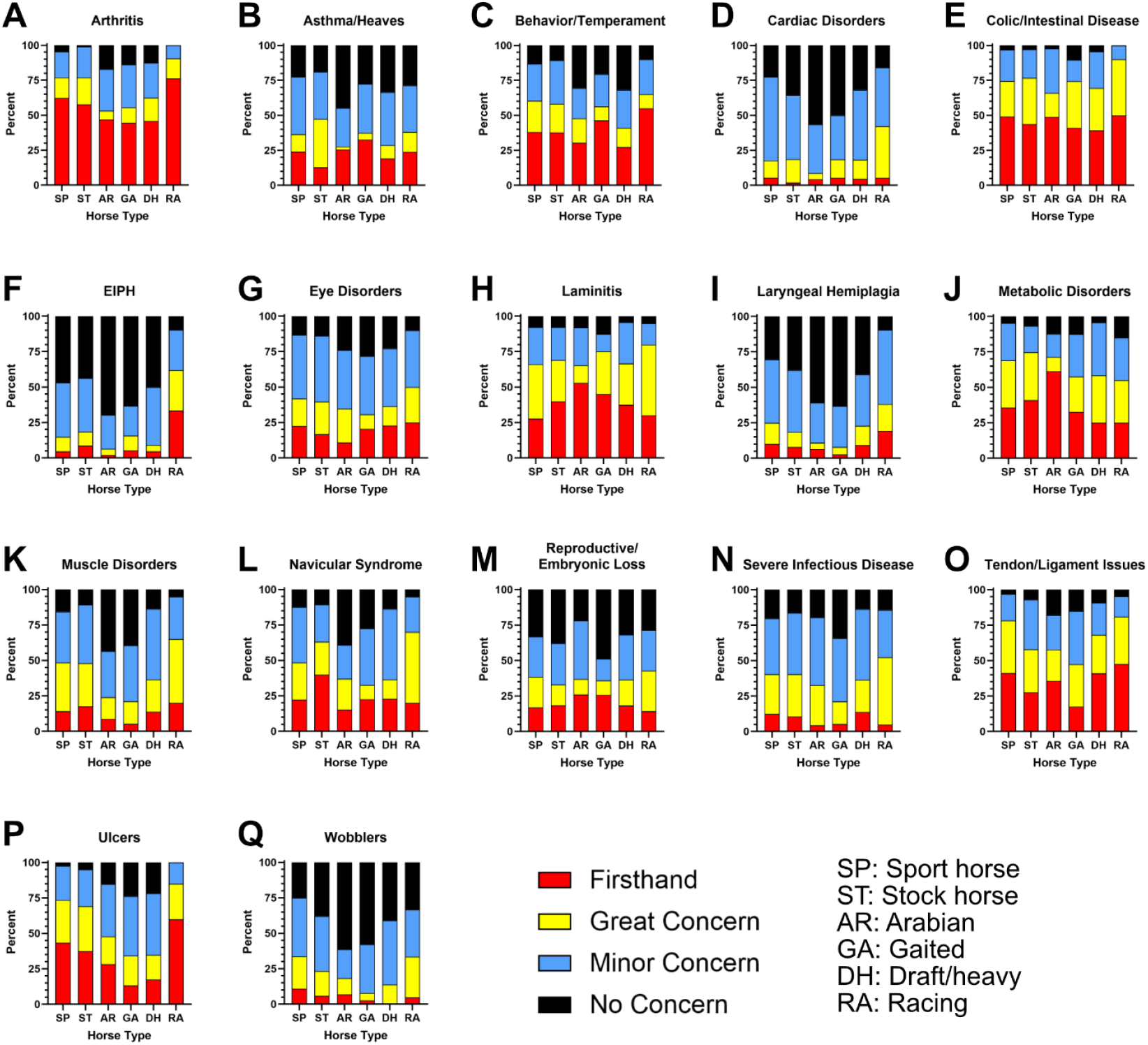
Prioritizing Herd Health Concerns. Survey respondents were asked about their level of concern about and/or firsthand experience with various herd health issues. Each panel shows the distribution of concern for those responding within each use category. Herd health issues queried include: (A) arthritis, (B) asthma or heaves, (C) behavior and temperament, (D) cardiac disorders, (E) colic and intestinal disease, (F) exercise-induced pulmonary hemorrhage (EIPH), (G) eye disorders, (H) laminitis, (I) laryngeal hemiplegia, (J) metabolic disorders, (K) muscle disorders, (L) navicular syndrome, (M) reproductive and embryonic loss issues, (N) severe infectious disease, (O) tendon and ligament issues, (P) ulcers, and (Q) wobblers or cervical vertebral malformation.

Those working with stock horses also represented nearly one-third of all respondents. Those responding indicated that they owned Quarter Horses (80.0%), Paint Horses (40.0%), and Appaloosas (9.2%), with many respondents owning horses of more than one breed within this category **(Table S2)**. Of the listed known genetic conditions of stock horses, respondents were most interested in learning more about PSSM, HYPP, and HERDA, among other diseases **(Figure S1)**. When asked about the level of concern for various herd health issues, these respondents prioritized the following concerns: arthritis, behavior and temperament, colic and intestinal disease, laminitis, metabolic disorders, navicular syndrome, tendon and ligament issues, and ulcers **(Figure 6)**. Stock horse responders most often identified PPID, anhidrosis, “kissing spines,” Type 2 polysaccharide storage myopathy (PSSM2), and OCD as being conditions of concern that they suspected were heritable **(Figure S2B)**.

For racing horses, respondents were most likely to work with Thoroughbreds (40.9%), with Standardbreds (27.3%) and Quarter Horses (27.3%) also represented **(Table S3)**. When asked about concern for various herd health issues, their prioritized concerns were the most numerous. They included arthritis, behavior and temperament, colic and intestinal disease, exercise-induced pulmonary hemorrhage (EIPH), laminitis, metabolic disorders, muscle disorders, navicular syndrome, severe infectious disease, tendon and ligament issues, and ulcers **(Figure 6)**. A few stakeholders (n = 8) answered the free-response question on conditions that stakeholders thought were genetic, with the top response being EIPH **(Figure S2C)**.

For Arabians, gaited horses, and draft and heavy breeds, only concern about herd health issues was solicited **(Figure 6)**. Prioritized concerns for Arabians included arthritis, colic and intestinal disease, laminitis, metabolic disorders, and tendon and ligament issues. Gaited horse concerns included arthritis, behavior and temperament, colic and intestinal disease, laminitis, and metabolic disorders. Draft and heavy horse breed concerns included arthritis, colic and intestinal disease, laminitis, metabolic disorders, and tendon and ligament issues.

## Discussion

To understand the genetic concerns of equine industry stakeholders in the United States as well as their knowledge levels of genetics and genetic testing, members of the USDA multistate project S1094 sought feedback via its first needs assessment offered online via Qualtrics in the spring and summer of 2024. Survey respondents were primarily owners and breeders, who represent the targeted demographic for the assessment. Findings from the survey revealed interesting information about the participating stakeholders, their level of confidence in their genetic knowledge, their greatest priorities based on concerns about herd health issues, and which conditions they suspect might be inherited.

A plurality of stakeholders responding to the survey were between the ages of 30 and 50 years old. This is similar to the median age (39 years old) of horse owners responding to the American Horse Council’s economic impact study of the horse industry in the U.S. (American Horse Council Foundation, 2023). Most respondents identified as horse owners in our survey; this differs significantly from the structure of the equine industry where approximately 1% of households in the U.S. own horses while just over 29% of households participate in or spectate at horse events without owning a horse (American Horse Council Foundation, 2023). Thus, horse owners were more likely to receive and/or complete our survey than other horse enthusiasts or industry members. Considering that many of the objectives of the multistate project are related to improving genetic testing resources, there is a benefit to reaching horse owners who are more likely to directly inquire about and access genetic testing for their horses.

In our survey, the two leading use categories for horses were sport horses and stock horses, comprising more than 63% of all respondents. Warmblood owners were the greatest percentage of sport horse respondents, despite only representing about 3% of horses reported in the impact study of the American Horse Council (American Horse Council Foundation, 2023). Thus, while representing a small percentage of horses in the United States, according to our survey, warmblood horse owners represent a vocal group of stakeholders with interest in genetic testing and concerns over genetic diseases in their breeds. The popularity of stock horses among the respondents was expected considering nearly half of all horses in the United States come from stock horse breeds (American Horse Council Foundation, 2023). Consequently, these two use categories represent the majority of stakeholder respondents in our survey with concerns about genetic testing and related genetic testing needs. Certainly, many genetic tests are currently commercially available for sport and stock horses (Bellone and Avila, 2020). Warmblood and stock-type breed registries and associations often fund genetic research studies that lead to genetic test development, aiding owners in breeding and management decisions.

Respondents from the racing horse use category were represented in smaller proportions (5.1%) than expected when considering the importance of racehorses to the equine industry. In 2022, direct and indirect impacts of the racing industry represented about $36 billion (American Horse Council Foundation, 2023). The limited number of responses represents a challenge for interpretation, suggesting that increased efforts to reach stakeholders in the racing industry are necessary for researchers in the multistate project. Still, these stakeholders, including statewide, regional, and national racing associations and foundations, have also invested in funding genetics research to resolve genetic issues in the breeds associated with racing.

When we asked respondents to consider their knowledge of genetics, the average respondent felt “confident” in their knowledge level. Thus, those responding to the survey likely represented people who are interested in genetics. This is useful information when considering respondents’ concerns about known genetic diseases and herd health issues. Most respondents indicated that they relied upon breed and discipline organizations as well as universities as resources for their knowledge of genetics. The predominant use of these two resources is logical considering that many breed and discipline organizations require genetic testing and most genetic tests to date have originated from research performed by equine geneticists at universities. Secondarily, respondents also relied upon horse owner community social media and veterinarians as resources for genetic information; these two resources are important for communicating equine genetic testing information (Hammons et al., 2020). These may represent an opportunity for the horse genetics research community to reach individual horse owners and caretakers directly with information about ongoing research, case recruitment, new tests, and other pertinent information. Further educational efforts of equine veterinarians in genetics and genomics could also support the overarching goal of disseminating knowledge to the industry, as they function as de facto educators.

Those participating in the survey were also generally interested in genetic testing strategies that could inform them about genetic diversity and performance genes in their horses. With improved familiarity of improvements in genomics resources and genetic testing developments, horse owners are more likely to use genetic testing Hammons et al., 2020). Maintaining unique breed characteristics often involves a degree of inbreeding as sires with desirable traits gain popularity. This can motivate breeders to weigh strategies for preserving genetic diversity while selecting for optimal traits. Genetic diversity has been studied recently in multiple breeds using several forms of genomic analyses to better understand how breeds have changed under the pressure of artificial selection for traits (Petersen et al., 2013; Cosgrove et al., 2020; Bailey et al., 2022; Esdaile et al., 2022; Bailey et al., 2024). Moreover, in 2010, with the discovery of variants in the myostatin gene *(MSTN)*, and then again in 2012 with the discovery of the “gait keeper” mutation in the doublesex and Mab-3 related transcription factor 3 gene (*DMRT3*), many stakeholders in the racing industry began to recognize the potential applications of analysis of equine performance genes and their variants (Hill et al., 2010; Andersson et al., 2012). With advances in genomic analysis and improvements in reference genomes (e.g., pangenomes, breed-specific reference genomes), the equine industry is poised to make advances in analysis of performance genes for stakeholders’ horses to improve their health, wellbeing, and performance (Finno and Bannasch, 2014; Avila et al., 2018; Bellone and Avila, 2020; Schaefer and McCue, 2020).

Respondents had many concerns about equine diseases and their heritability potential. Since causative mutations for diseases like HYPP were first discovered, horse owners have come to utilize genetic testing for breeding and management decisions (Rudolph et al., 1992). While HYPP and other similar conditions for which genetic tests already exist are simple genetic diseases, controlled by a mutation in a single gene, many of the diseases and conditions of concern identified by respondents across breed/discipline categories are complex, likely involving many genes and environmental influences. These included arthritis, colic and intestinal disease, laminitis, metabolic disorders, tendon and ligament issues, navicular syndrome, gastric ulcers, and muscle disorders. Anhidrosis and overriding dorsal spinous processes were also commonly identified as suspected heritable conditions not provided on the pre-populated list. Furthermore, respondents were concerned with behavior and temperament issues, which are also considered quantitative, complex traits (Wickens and Brooks, 2020). While research into genetic risk factors for various complex diseases is ongoing, collaborative efforts will likely be required to thoroughly explore the role of specific genes and variants in each of these diseases to determine if development of valid genetic tests is even possible (McCoy et al., 2019; Norton et al., 2019; Petersen et al., 2019; Kingsley et al., 2022; Kapusniak et al., 2025). A goal of the multistate research project S1094 is to bring together teams of geneticists across the country to recruit large sample populations, thoroughly scrutinize horses’ phenotypes, and combine unique sets of skills amongst researchers to unravel genetic mechanisms underlying these complex diseases and conditions.

Overall, survey respondents recognized the impact that genetics and genetic testing have on the equine industry. Through improved understanding and judicious use of genetic testing, owners can make more informed breeding decisions that produce healthier and better performing horses. Moreover, healthier horses generally live longer, have fewer costly conditions to manage, and may be more efficient at breeding. Furthermore, realizing that selection of some traits might promote deleterious phenotypes, with robust genetic testing strategies, breeders and owners can carefully weigh their breeding options and can be better prepared for future management issues of those progeny.

Needs assessments from stakeholders in the equine industry are indispensable to the researchers in the S1094 multistate project. From this survey, we determined that horse owners were strongly interested in genetics. We also learned that most survey respondents felt relatively confident in their understanding of horse genetics, and we determined from where they were drawing their knowledge of genetics. Moreover, responding stakeholders indicated that they were interested in evaluating genetic diversity and performance traits, and we were given insight into a myriad of concerns about herd health issues. Many of those diseases and conditions are quite complex; thus, the participants in the S1094 multistate project will benefit from working together to develop new genetic testing tools. There is also a need for novel educational materials to be distributed through a variety of outlets to help horse owners make informed breeding decisions and determine how future offspring will be managed given what is known about their genetic composition.

## Abbreviations

AI: artificial intelligence
AR: Arabian horses
CVSM: cervical vertebral stenotic myelopathy
DH: draft and heavy horses
DMRT3: doublesex and Mab-3 related transcription factor 3 gene
DSLD: degenerative suspensory ligament desmitis
EIPH: exercise-induced pulmonary hemorrhage
FFS: fragile foal syndrome
GA: gaited horses
GBED: glycogen branching enzyme deficiency
HERDA: hereditary equine regional dermal asthenia
HYPP: hyperkalemic periodic paralysis
IMM: immune mediated myositis or myosin-heavy chain myopathy
LWO: lethal white overo syndrome
MH: malignant hyperthermia
MSTN: myostatin gene
OCD: osteochondritis dissecans
ORDSP: overriding dorsal spinous processes
PSSM: polysaccharide storage myopathy
PPID: pituitary pars intermedia dysfunction, or Cushing’s Disease
RA: racing horses
RLN: recurrent laryngeal neuropathy
SP: sport horses
ST: stock horses
USDA: United States Department of Agriculture

## Acknowledgements

For their help with testing the survey, we would like to acknowledge Ashley Griffin of the Extension Foundation, Dr. Craig Wood form the University of Kentucky, Jessica Hein from the American Paint Horse Association, and Dr. Neely Walker from Louisiana State University.

## Funding

Efforts of M.J.M. are supported by University of California Division of Agricultural and Natural Resources (UC ANR) through its Agricultural Experiment Station at UC Davis (CA-D-ASC-2760-RR). This research was supported in part by the intramural research program of the U.S. Department of Agriculture, National Institute of Food and Agriculture, Hatch Project [7001788].

## Conflict of interest statement

SH, EN, LPR, EMN, AMM, and CW have no conflicts. MJM’s spouse directs the University of California Davis Veterinary Genetics Laboratory; their spouse was not involved in the design, execution, or interpretation of the study. SAB advises Etalon Diagnostics. MEM receives income from PSSM1 genetic test sales; their university manages royalties received from genetic tests developed.

## Author contributions

Michael J. Mienaltowski (Conceptualization, Survey methodology, Survey respondent recruitment, Data curation, Formal analysis, Writing—original draft, Writing—review & editing), Sarah Hernandez (Data curation, Formal analysis, Writing—original draft, Writing—review & editing), Emma Nastrini (Data curation, Formal analysis, Writing—original draft, Writing—review & editing), Carissa L. Wickens (Survey initialization, Survey methodology, Survey respondent recruitment, Writing—review & editing), Molly E. McCue (Survey respondent recruitment, Writing—review & editing), Laura Patterson Rosa (Survey respondent recruitment, Writing— review & editing), Elaine M. Norton (Conceptualization, Survey methodology, Survey respondent recruitment, Writing—review & editing), Annette M. McCoy (Conceptualization, Survey methodology, Survey respondent recruitment, Writing—review & editing), and Samantha A. Brooks (Conceptualization, Survey initialization, Survey methodology, Survey respondent recruitment, Writing—review & editing)

## Notes

### Competing Interest Statement

SH, EN, LPR, EMN, AMM, and CW have no conflicts. The spouse of MJM directs a university veterinary genetics diagnostic laboratory, but their spouse was not involved in the design, execution, or interpretation of the study. SAB advises Etalon Diagnostics. MEM receives income from PSSM1 genetic test sales, and their university manages royalties received from genetic tests developed.

