## Supplemental Information for "United States stakeholder insights on genetic testing for equine health and breeding"

**Table S1. Sport horse breeds represented.**

| Breed | Number | Percent |
|---|---|---|
| Warmblood | 88 | 68.2% |
| Thoroughbred | 25 | 19.4% |
| Quarter Horse | 6 | 4.7% |
| Connemara ponies | 2 | 1.6% |
| Welsh cob | 2 | 1.6% |
| Other | 6 | 4.7% |

**Table S2. Stock horse breeds represented.**

| Breed | Number | Percent |
|---|---|---|
| Quarter Horse | 96 | 80.0% |
| Paint | 48 | 40.0% |
| Appaloosa | 11 | 9.2% |
| Mustang | 3 | 2.5% |
| Appendix | 2 | 1.7% |
| Other | 19 | 15.8% |

“Percent” listed represents the percentage of stock horse respondents indicating that they own horses of that breed. Respondents could own more than one breed.

**Table S3. Racing horse breeds represented.**

| Breed | Number | Percent |
|---|---|---|
| Thoroughbred | 9 | 40.9% |
| Standardbred | 6 | 27.3% |
| Quarter Horse | 6 | 27.3% |
| None Listed | 1 | 4.6% |

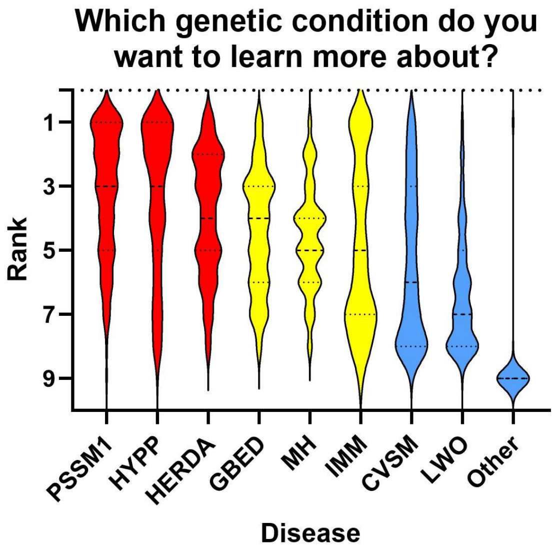


**Figure S1. Genetic concerns stock horse owners want to learn about.** Violin plots represent distributions of ranks assigned for each of the conditions from 1 (most concern) to 9 (least concern). Conditions were PSSM1 (polysaccharide storage myopathy 1), HYPP (hyperkalemic periodic paralysis), HERDA (hereditary equine regional dermal asthenia), GBED (glycogen branching enzyme deficiency), MH (malignant hyperthermia), IMM (immune mediated myositis or myosin-heavy chain myopathy), CVSM (cervical vertebral stenotic myelopathy), and LWO (lethal white overo syndrome).

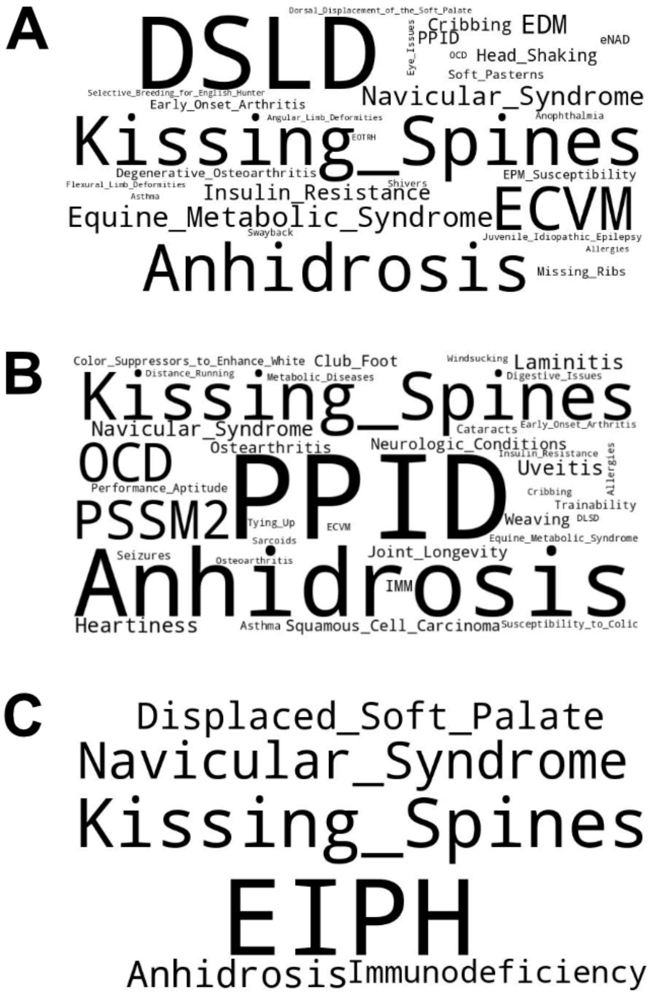


**Figure S2. Word clouds of respondents' suggestions for heritable conditions.** Respondents were asked about which conditions they felt might be heritable. Responses for **(A)** sport horses, **(B)** stock horses, and **(C)** racing horses are provided in word clouds, which were generated with Copilot AI given instructions to make text size proportional with the number of mentions in survey responses. The number of responses offered were n= 77 for sport horses, n = 62 for stock horses, and n = 8 for racing horses.
